# Determinants of haemosporidian single and co-infections risks in western paleartic birds

**DOI:** 10.1101/2021.06.11.448006

**Authors:** Romain Pigeault, Mathieu Chevalier, Camille-Sophie Cozzarolo, Molly Baur, Mathilde Arlettaz, Alice Cibois, André Keiser, Antoine Guisan, Philippe Christe, Olivier Glaizot

## Abstract

Co-infections with multiple pathogens are common in the wild and may act as a strong selective pressure on both host and parasite evolution. Yet, contrary to single infection, the factors that shape co-infection risk are largely under-investigated. Here, we explored the extent to which bird ecology and phylogeny impact single and co-infection probabilities by haemosporidian parasites using large datasets from museum collections and a Bayesian phylogenetic modelling framework. While both phylogeny and species attributes (e.g. size of the geographic range, life-history strategy, migration) were relevant predictors of co-infection risk, these factors were less pertinent in predicting the probability of being single infected. Our study suggests that co-infection risk is under a stronger deterministic control than single-infection risk. These results underscore the combined influence of host evolutionary history and species attributes in determining single and co-infection pattern providing new avenues regarding our ability to predict infection risk in the wild.

## INTRODUCTION

One of the most fundamental questions regarding the evolutionary ecology of host–parasite interactions consists in identifying the factors shaping infection risk and parasite distribution among hosts (Sutherland *et al.* 2013). Beyond identifying the most at-risk species in a given community (Greenberg *et al.* 2017), studying the ecological and the evolutionary factors that drive heterogeneity in hosts’ infection risk at both the intra- and interspecific levels is paramount to understand the emergence, dynamic and evolution of infectious diseases (Rodrigues *et al.* 2009; Streicker *et al.* 2013).

A number of studies have attempted to identify the factors involved in infection risks (e.g. Ricklefs 1992; Greenberg *et al.* 2017; Barrow *et al.* 2019; Ellis *et al.* 2020). At the intraspecific level, variation in infection risks has been associated to several host-specific features related to parasite exposure (e.g. individual differences in behavior, Ezenwa *et al.* 2016 and/or parasite susceptibility (e.g. body condition, Beldomenico *et al.* 2008; sex, Christe *et al.* 2007; age, Channappanavar & Perlman 2020, or reproductive stages, Christe *et al.* 2000). At the interspecific level, several studies have shown that the susceptibility to parasites may be conserved through host phylogeny (e.g. Hoverman *et al.* 2011; Greenberg *et al.* 2017; Barrow *et al.* 2019). This phylogenetic signal can be explained by the inheritance of parasites from a common ancestor (Davies & Pedersen 2008) or by the role played by different evolutionarily conserved factors (e.g. genetic characteristics of the immune system,Minias *et al.* 2019; host physiological phenotype, Cronin *et al.* 2010; geographic range size, Waldron 2007; life-history strategies, Valenzuela-Sánchez *et al.* 2021). Phylogenetically distant species can nevertheless present similar infection risk patterns if they share ecological traits that have an influence on the exposure or the susceptibility to parasites (e.g. behavior, Han *et al.* 2015; foraging and living environment, Menzies *et al.* 2021).

In nature, host individuals encounter a multitude of different parasites during their lifetime. Simultaneous infections by multiple pathogen strains or species are therefore very common (Read & Taylor 2001; Bordes & Morand 2011; Hoarau *et al.* 2020). Similar to single infections, co-infections (also termed multiple infections or polyparasitism) are ubiquitous and are heterogeneously distributed among hosts (Viney & Graham 2013). Indeed, the probability of being co-infected by two or more parasites has been shown to vary considerably at both the intra-(Brooker & Clements 2009; Susi *et al.* 2015) and interspecific (Thurber *et al.* 2013; González-González *et al.* 2014) levels. However, while the drivers of single infections (or infections in general) are well studied, the drivers of co-infection risks are currently largely understudied, co-infection being either ignored (e.g. Soares *et al.* 2020; Starkloff *et al.* 2020) or not mentioned (e.g. Barrow *et al.* 2019; de Angeli Dutra *et al.* 2021). Yet, co-infections are of paramount importance for understanding the ecology and evolution of host-parasite interactions owing to the wide range of effects they have on host fitness (e.g. Marzal *et al.* 2008; Bordes & Morand 2011). They are however difficult to predict because the cumulative effects resulting from the interaction between parasites can be different from the sum of individual effects (“one plus one is not two“; Marzal *et al.* 2008). Beyond the direct consequences on host fitness, co-infections may also influence the evolutionary trajectories of pathogen populations through their effects on the evolution of virulence (Alizon *et al.* 2013) and therefore on the within-host infection dynamic (Susi *et al.* 2015). Likewise, interactions between co-infecting parasites may contribute to the maintenance of the genetic variation of each parasite and impact host-parasite co-evolution (Seppälä & Jokela 2016). Altogether, these results suggest that co-infections are important drivers of both parasite evolution and epidemiology (Tollenaere *et al.* 2016).

Individual risk of infection is the product of the host’s exposure and susceptibility to parasites. This exposure-susceptibility interaction becomes more complex as the number of parasites potentially involved in the association increases (Viney & Graham 2013). At first glance, one can assume that evolutionary histories and species attributes affecting single infection probabilities would affect co-infection probabilities in a similar way. However, some attributes may be preponderant for co-infections. For instance, species with large geographic ranges and/or climatic niches are more likely to be found in habitats that are suitable for different parasites and are therefore more likely to encounter (and thus to be infected by) multiple parasites. Similarly, migration may bring hosts into contact with a larger diversity of parasites (Smith *et al.* 2004; Hellgren *et al.* 2007). The position of species along the slow-fast life-history continuum could also be a strong predictor of co-infection risk (Vaumourin *et al.* 2015; Valenzuela‐Sánchez *et al.* 2021). For instance, the differential allocation of resources to immunity between fast- and slow-living species may affect susceptibilities to parasite infection (Valenzuela‐Sánchez *et al.* 2021) and ultimately impact the frequency of co-infections (Vaumourin *et al.* 2015).

Avian haemosporidian parasites (phylum Apicomplexa: order Haemosporida: Family: Plasmodiidae: *Plasmodium*, *Haemoproteus* and *Leucocytozoon*, (Valkiunas 2004) offer an exciting opportunity to better understand the distribution of co-infections and the relative role of both ecological and phylogenetic factors. These blood parasites, transmitted by hematophagous dipteran insects, are distributed worldwide and are found in most bird families (Valkiunas 2004). Co-infections by different haemosporidian parasite genera have been shown to predominate in some avian populations (e.g. Marzal *et al.* 2008; Clark *et al.* 2016; Pigeault *et al.* 2018). Although few studies have evaluated the impact of co-infection by two haemosporidian parasites on host fitness, some evidence suggest that they may negatively impact bird survival rate (Marzal *et al.* 2008; Palinauskas *et al.* 2011; Pigeault *et al.* 2018), thus inducing a selective pressure on host evolution.

Here, we quantified for the first time the impact of multiple host ecological attributes and phylogeny, on both single and co-infection risks in 151 European bird species. Specifically, we used phylogenetic mixed-effect models to estimate the impact of climatic niche breadth and position, geographic range size, trophic niche characteristics, life-history strategies and behavioral characteristics (nesting and migration behaviors) on the probability to be single or co-infected by haemosporidian parasites. Bird infection status was estimated by molecular methods using tissue collections from the museums of Lausanne and Geneva collected on dead individuals found mainly in western Switzerland. Our results confirm that co-infections are as widespread as single infections and that both are heterogeneously distributed among species. We further confirm the role of species attributes and phylogeny in predicting infection status. But most importantly, we show that our ability to predict co-infection risk is much larger than for single infections, suggesting that deterministic processes related to evolutionary history and ecological characteristics may be more important for the former than for the latter. Our results provide new insights regarding our ability to predict infection risk and the potential role played by co-infections in driving hosts evolution.

## MATERIAL AND METHODS

### Avian samples and parasites detection

Our data set includes 1361 samples of 151 species encompassing 44 families and 18 orders (**Appendix 1**, **Table S1**). Sampling was conducted on salvaged birds that were obtained between 1990 and 2019. and consisted of tissues (muscle and liver) stored in 85% ethanol at 4°C at the Cantonal Museum of Zoology in Lausanne (855 specimens) and in 90% ethanol at - 20°C at the Natural History Museum of Geneva (506 specimens).

For each individual, parasites (i.e. *Haemoproteus, Leucocytozoon* and *Plasmodium*) were detected from tissue samples using molecular methods. Specifically, a nested PCR (Hellgren *et al.* 2004) was performed in triplicates on all samples after DNA was extracted from tissues using a DNeasy Blood & Tissue Kit (Qiagen, Switzerland) following the manufacturer’s instructions. Nested PCR products were visualized on agarose gels after electrophoresis to identify infected samples. This nested PCR protocol does not allow to detect co-infections between parasites of the genera *Haemoproteus* and *Plasmodium*. Therefore, all positive samples were sequenced in both directions as in van Rooyen *et al.* (2013) and identified by performing a local BLAST search in the MalAvi database (http://mbio-serv2.mbioekol.lu.se/Malavi/; Bensch *et al.* 2009). Co-infections by *Haemoproteus* and *Plasmodium* were identified by analyzing double nucleotide peaks on sequence chromatographs. We re-amplified and re-sequenced all the samples for which the chromatograph could not reliably identify the parasite sequences. All sequences were edited using Geneious v8.0.5. Birds not infected by any parasites were classified as “not infected”, birds infected with a single parasite genus were classified as “single infected” and those infected with at least two different genera as “co-infected” (Pigeault *et al.* 2018).

### Species attributes

#### (a) Life-history strategies, ecological and behavioral characteristics

We used published trait data to position each species along the slow-fast continuum of life-history variation (Storchová & Hořák 2018). More specifically, bird position was represented by the first axis of a principal component analysis (PCA) performed on nine variables describing bird reproductive traits (clutch size, number of broods per year, average length, width and weight of the egg, incubation period, fledging age, age at first breeding) and maximum lifespan (**Appendix 1, Table S1**). The first axis explained 62.7% of the variability and represented a gradient going from fast (negative values) to slow (positive values) life-history strategies.

The trophic niche of bird species was estimated using 35 variables describing the diet during the breeding season (Pearman *et al.* 2014). Specifically, we considered 14 variables characterizing diet (*e.g.* seeds, fruits, invertebrates, fishes), nine variables characterizing food acquisition behavior (*e.g.* air pursuit, foliage glean, dig, probe), nine variables characterizing the substrate from which food is taken (*e.g.* air, water surface, mud, canopy) and three variables characterizing the daily foraging period (**Appendix 1**, **Table S2**). As in Pearman *et al.* (2014), we also included body weight as a surrogate for total energy requirements. These variables were scored as either 0 or 1, with the exception of body weight, which was scored as the average weight of individuals during the breeding season (Pearman *et al.* 2014). Trophic niches were represented by the scores of each species along the first two axes of a Hill-Smith ordination (denoted OA; Hill & Smith 1976). These axes roughly corresponded to the structure (from open to forest habitats; OA1 = 19.3%) and the height (from underwater and ground to foraging in trees or in flight; OA2 = 12.6%) of the foraging environment.

The remaining traits (nest type and migration status) were extracted from Storchová & Hořák (2018). Nest type was categorized as either “open” or “closed” while migration status was categorized as “sedentary” (species living in the same area in both the breeding and the non-breeding season), “migratory” (species migrates between breeding and non-breeding season) and “facultative migrant” (species makes irregular shifts in breeding and/or nonbreeding season).

#### (b) Climatic niche breadth, climatic niche position and geographic range size

Estimating species climatic realized niches (i.e. the set of suitable environmental conditions accessible to the species and constrained by biotic interactions; Jackson and Overpeck (2000)) requires data for the full geographical range of species together with the corresponding environmental variables (Guisan *et al.*, 2017). For each species, we estimated its climatic niche by cross-referencing IUCN range maps (https://www.iucnredlist.org/; only considering the resident and the breeding range) with climatic data. Climatic layers for 19 bioclimatic variables were extracted from worldclim (https://www.worldclim.org/) at a 10’ resolution (roughly 340 km² at the equator). From these variables, we performed a PCA from which we extracted the two first axes, which explained 55% and 19% of the total variance, respectively. Owing to the difficulty to characterize the climatic niche of migratory species, analyses presented below were repeated with full and facultative migrants excluded (see **Fig. S1**, **Fig. S2B**).

To obtain the climatic niche of each species, we first projected the environmental values corresponding to the geographic range coordinates falling inside the IUCN polygon within the two-dimensional space defined by the two PCA axes. We then used a kernel density estimator (KDE) to delineate species envelopes (**Fig. 1**). KDE have proved useful to characterize complex and potentially irregular shapes (Blonder *et al.* 2014) and are increasingly used to characterize climatic niches (e.g. Broennimann *et al.* 2012). For each species, the bandwidth of the KDE was estimated from the data using a Hpi multivariate generalization of the plug-in bandwidth selector (Wand & Jones 1994). Envelopes were then defined as the minimum threshold of probability density that included 99% of points (to leave out environmentally atypical occurrences). From species envelopes, we extracted its area (niche breadth) and computed its centroid as the mean of point coordinates falling inside the delimited niche. We then extracted the coordinates of the centroid on each of the two PCA axes (**Fig. 1**). To test the robustness of our results we used two other algorithms to delineate species realized niches: convex hulls and alpha hulls (see **Fig. S2**, **Fig. S3**). The area of the geographic range was directly extracted from IUCN polygons.

**Figure 1.**
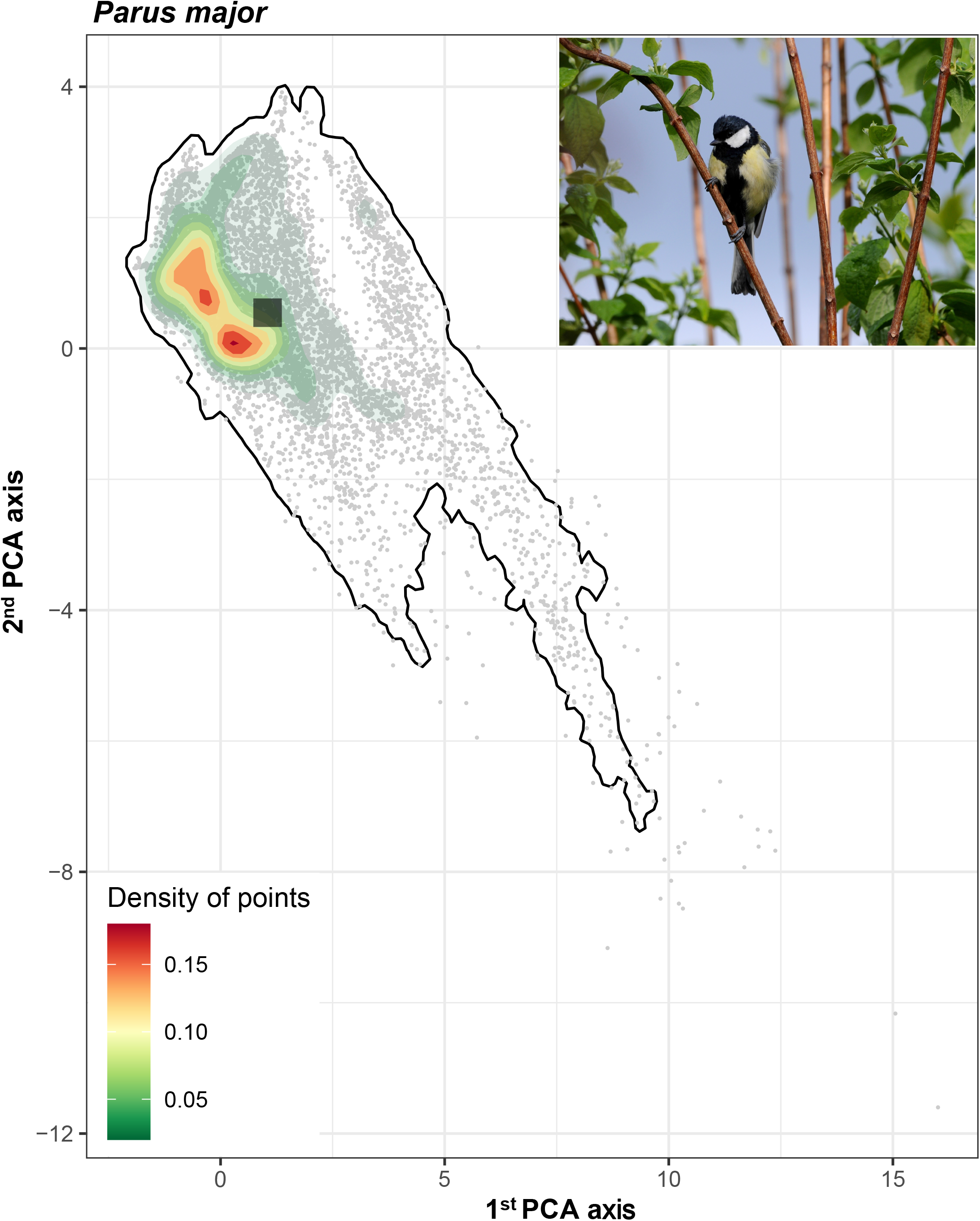
Representation of the climatic niche of *Parus major*. Species climatic niche represented in the two-dimensional space defined by the two first PCA axes performed on a set of 19 bioclimatic variables. The envelope was estimated using a Kernel Density Estimator including 99% of occurrence point (in grey) while the centroid was computed as the average coordinates of occurrences (The picture of *Parus major* was taken by *Philippe Christe* ©).

### Infection prevalence among avian phylogenetic tree

To illustrate the effects of host evolutionary history on single and co-infection probabilities, we followed the methods of Barrow *et al.* (2019). Specifically, we used 1000 trees from BirdTree.org (backbone tree from Hackett *et al.* 2008) to generate a consensus tree using the *ls.consensus* function (package phytools v. 0.6-00, Revell 2020). The prevalence of infections (i.e. the proportion of birds infected by haemosporidian parasites altogether and by each haemosporidian genus separately) was calculated for each species and mapped on the phylogenetic tree using the *contMap* function (package phytools). The prevalence of infection was visualized over the whole phylogeny using species for which there were at least 5 samples.

### Statistical analyses

Overall, we computed ten predictors: the first PCA axis summarizing bird life-history strategies (i.e. position along the slow-fast life history continuum), the two OA axes summarizing bird trophic niches, the climatic niche breadth, the geographic range size, the coordinates of the climatic niche centroid on each PCA axis, the migration status, the nest type and the museum affiliation (to account for potential effect related to tissue collection and/or conservation). To prevent collinearity between predictors, we checked that variables had Pearson’s correlation coefficients ρ ≤ |0.7| (Dormann *et al.* 2013). We found that OA1 and the geographic range size were strongly correlated with bird life-history strategies (ρ = −0.81) and the climatic niche breadth (ρ = 0.76), respectively. These variables were thus removed from the analysis. However, we also tested the effect of these variables in separate models where they replaced the variables they were correlated with.

We built two types of phylogenetic multilevel models using Bayesian multilevel models from the *brms* package (Bürkner *et al.* 2021). First, we used a phylogenetic multinomial model with a “Categorical” error distribution to study the effects of species attributes and phylogeny on the probability to be single and co-infected. Second, we used three phylogenetic generalized linear multilevel models with a “Bernoulli” error distribution to study the effects of species attributes and phylogeny on the probability to be infected by each parasite genus, separately. For all models, host species and phylogeny were treated as random effects. Continuous predictors were standardized to z-scores (mean=0, variance=1) to improve model convergence and parameter estimation. We used the default priors of the *brms* package (i.e. weakly informative priors) and ran three chains with 11,000 iterations. The first 1,000 iterations were considered as burn-in and were thus discarded. Chains were thinned every 10 iterations. To account for phylogenetic uncertainty, we randomly sampled 100 trees from the set of trees extracted from BirdTree.org and ran the analyses on each tree. We then combined all models using the *combine_models* function. Note that this approach critically depends on the assumption that all trees are equally likely. We followed the methods of Barrow *et al.* (2019) to compute the phylogenetic signal of single and co-infection probabilities estimated from the multinomial model and of infection probability by each genus from the three generalized linear models.

## RESULTS

Unless otherwise stated, all results were consistent regardless of the method used to delineate species envelopes as well as when migratory species were removed.

### Single and co-infection prevalence

We detected 486 infected individuals (35.7%), among which 101 (7.4%) were co-infected by at least two different parasite genera (20.8% of infected birds). Focusing on species for which there were at least 5 samples, we observed a large variation in the prevalence of infections across the phylogeny (**Fig. 2**). Species with the highest prevalence (>90% of infected individuals) included *Asio otus*, *Falco subbuteo* and three passerines: *Corvus corone*, *Coccothraustes coccothraustes* and *Turdus philomelos* (**Fig. 2**). In contrast, eleven species belonging to eight different genera were never infected (**Fig. 2**). The rate of co-infection greatly varied between species, ranging from zero individual in 32 species to more than 50% individuals in seven species (**Fig. 2**). The most frequent parasite genus was *Leucocytozoon* (51.03% of infected birds) followed by *Plasmodium* (38.65%) and *Haemoproteus* (26.38%). The infection prevalence of each haemosporidian genus also varied greatly across the phylogeny (**Fig. S4**) with some families exhibiting higher infection prevalence than others. For instance, *Leucocytozoon* infection prevalence was relatively high for Corvidae (78%) whereas *Plasmodium* infection prevalence was the highest for Turdidae (86%).

**Figure 2.**
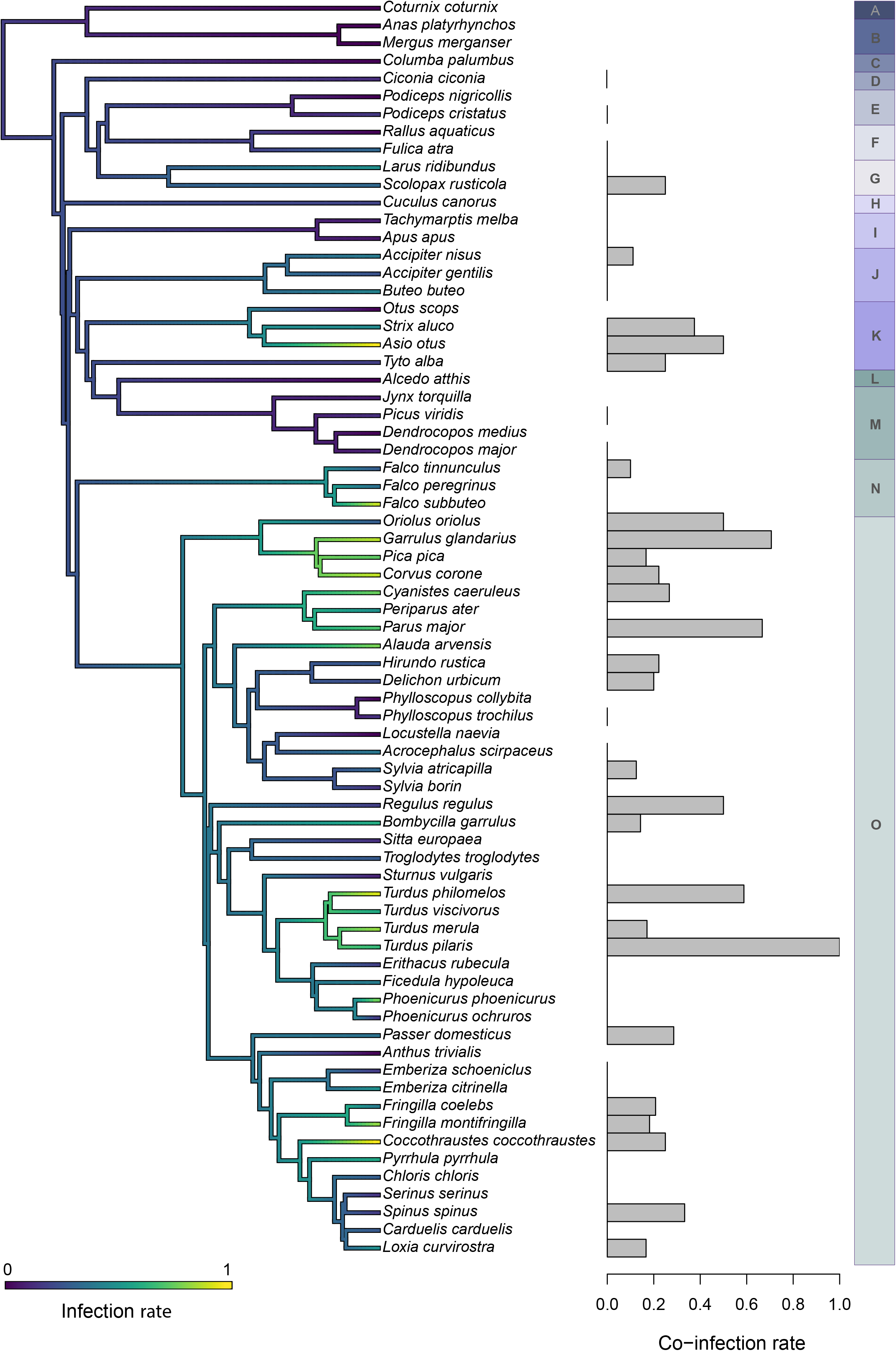
Haemosporidian infection across the avian phylogeny. The proportion of infected individuals (single and co-infected), for each bird species for which at least five individuals were sampled, was mapped as a continuous trait using the *contMap()* function in the *phytools* R package (Revell 2020). The consensus tree was generated from 1000 phylogenies obtained from BirdTree.org. The bar plot represents the proportion of co-infected individuals within the species with at least one infected individual. Each letter (from A to O) corresponds to a bird order. A: Galliforme, B: Anseriformes, C: Columbiformes, D: Ciconiiformes, E: Podicepediformes, F : Gruiformes, G : Charadriiformes, H : Cuculiformes, I : Apodiformes, J : Accipitriformes, K : Strigiformes, L : Coraciiformes, M : Piciformes, N : Falconiformes, O : Passeriformes

### Predictors of haemosporidian single and co-infection

The structure (trophic niche, OA1) and the height of the foraging environment (OA2), the area of the climatic niche (niche breadth) and the location of the niche centroid were not pertinent to explain the probability to be single or co-infected (i.e. the 95% confidence interval [CI] overlapped with 0, **Fig. 3**). We note however that while this absence of effect was generally robust to the type of algorithm used to delineate climatic niches (**Fig. S2**) and the removal of migratory species, we detected a tendency toward an effect of niche breadth on co-infection probability when excluding migratory species (**Fig. S1, Fig. S2**)). Migratory species (full and facultative migrants) and species with open nests presented a higher probability to be single and co-infected than species with the opposite characteristics (sedentary species and species with closed nests; **Fig. 3**). Further, we found that species life-history strategies and geographical range size influenced the co-infection probability with a higher probability for co-infection for fast-living species (**Fig. 3**) and for species distributed on a large geographic area (**Fig. 3, Fig. S5**). Both single and co-infection probabilities presented a strong evidence for a phylogenetic effect but the signal was much lower for single (0.12 [95% CI 0.02–0.26]) than for co-infections (0.53 [95% CI 0.25–0.76]).

**Figure 3.**
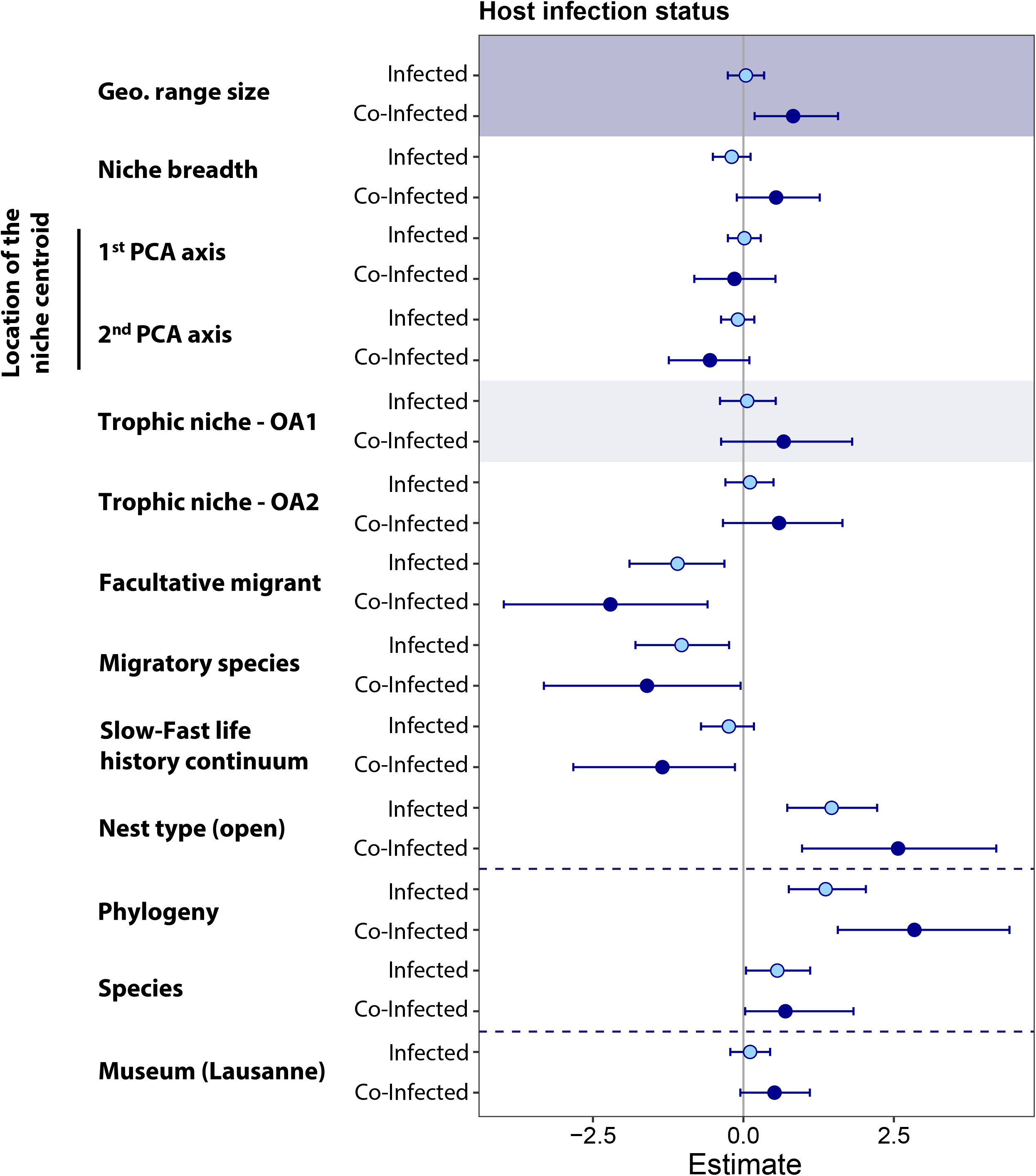
Posterior mean and associated 95% credible intervals of predictors and random effects (i.e. phylogeny and host species) as estimated by *brms* model with infection status (i.e. uninfected, infected, co-infected) as the response variable. Effects shown are relative to the uninfected reference category, with single infection in light blue and co-infection in dark blue. The posterior distribution of predictors and random effects with a negligible effect on infection and co-infection probabilities were expected to be centered on zero. For categorical predictors, the effects shown are relative to the reference categories: nest type (closed), migration behavior (resident), museum (Lausanne). Museum affiliation was added as a predictor to account for potential effect related to tissue collection and/or conservation. Since OA1 and the geographic range size were strongly correlated with bird life-history strategies (ρ = −0.81) and the climatic niche breadth (ρ = 0.76) the posterior mean of OA1 and geographic range size presented here (grey and blue rectangles) was estimated from two other *brms* model whose complete results are shown in **Fig. S5**.

Taken together, species attributes and phylogeny explained 50% [95% CI 43.3–55.0] of the variability in co-infections probability but only 26% [95% CI 21.4–29.7] of single infection probability. Furthermore, while effect sizes tended to go in the same direction, the magnitude of effects was much larger for co-infection than for single infection probability.

### Predictors of infection by each haemosporidian genus

None of the tested variables appeared to be a strong predictor of *Plasmodium* infection probability (**Fig. 4, Fig. S3**). On the other hand, *Haemoproteus* infection probability tended to be impacted by both the height of the foraging environment and species life-history strategies thus indicating a lower probability of infection for slow-living species and species foraging on the ground (**Fig. 4**). Species with open nests also tended to have higher *Haemoproteus* prevalence than species with closed nests (**Fig. 4**). Regarding *Leucocytozoon*, infection probability tended to increase with the geographic range size (**Fig. 4**) but to be lower for facultative migrants and species with closed nests (**Fig. 4**). The phylogenetic signal was the highest for *Leucocytozoon* (0.50 [95% CI 0.22–0.73]), followed by *Plasmodium* (0.33 [95% CI 0.10–0.61]) and then by *Haemoproteus* (0.24 [95% CI 0.02–0.59]).

**Figure 4.**
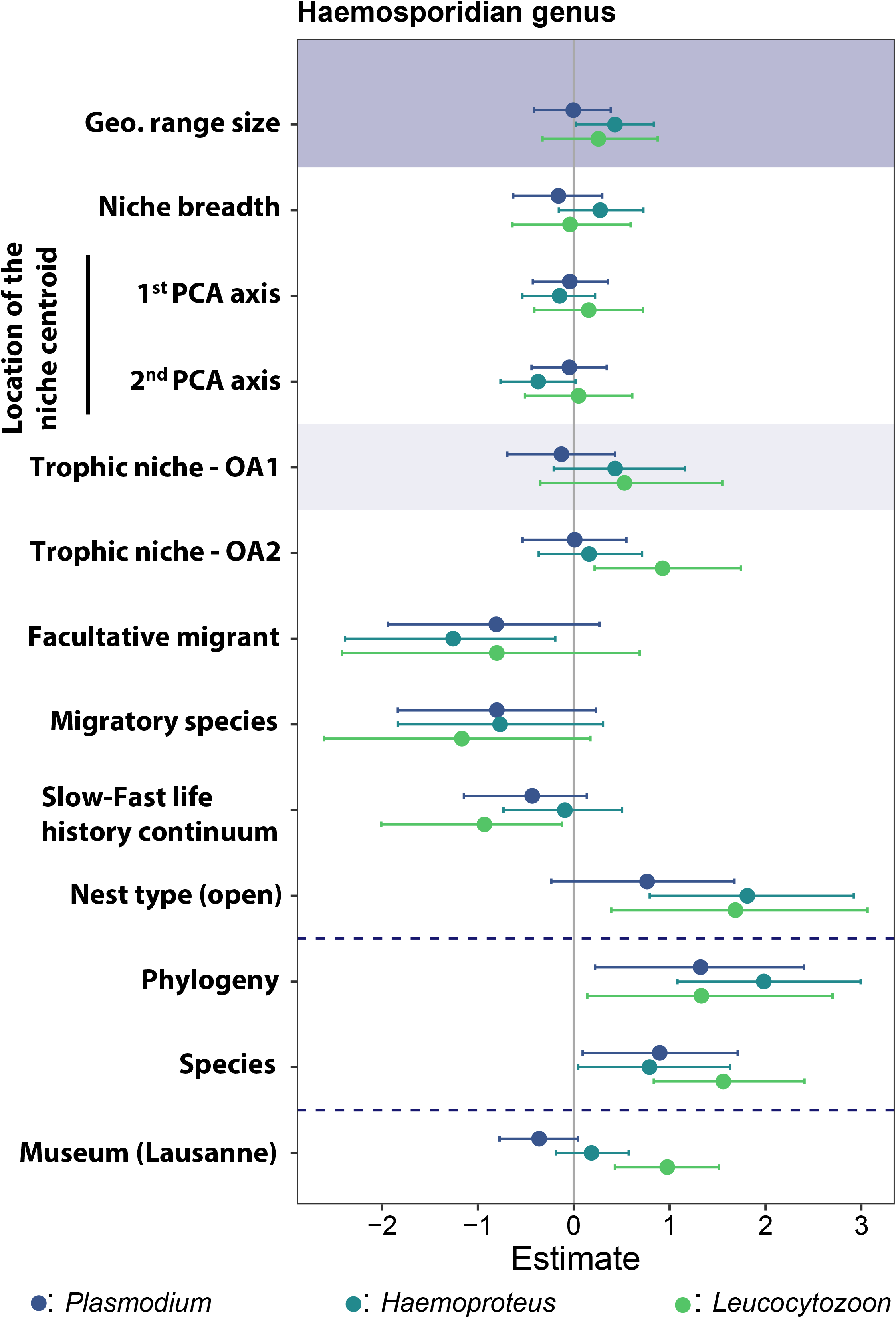
Posterior mean and associated 95% credible intervals of predictors and random effects (i.e. phylogeny and host species) as estimated by *brms* models with infection status (uninfected vs. infected) by each parasite genera as the response variable. For categorical predictors, the effects shown are relative to the reference categories: nest type (closed), migration behavior (resident), museum (Lausanne). Museum affiliation was added as a predictor to account for potential effect related to tissue collection and/or conservation. Since OA1 and the geographic range size were strongly correlated with bird life-history strategies (ρ = −0.81) and the climatic niche breadth (ρ = 0.76) the posterior mean of OA1 and geographic range size presented here (grey and blue rectangles) was estimated from two other *brms* model whose complete results are shown in **Fig. S6**.

## DISCUSSION

In this study, we evaluated the extent to which host phylogeny and species attributes related to climatic niche properties and other ecological and life-history traits influenced the probability of haemosporidian infection and co-infection in western Palearctic birds. We found that (1) while some attributes (e.g. migration) influenced both the probability of being single and co-infected, others (e.g. geographic range size) were only pertinent in explaining variation in co-infection probability and (2) that phylogeny is a far more important predictor of co-infection probability than of single infection probability. Overall, the effect size of all predictors and the proportion of variance explained by our models were significantly larger for co-infection than for single infection probability. Altogether, these findings suggest that co-infection probability is under a stronger deterministic control than single infection which may rather be influenced by stochastic processes (e.g. random encounter rate with parasites) or by other factors not included in our study (e.g. hosts’ features).

Two parameters associated with host ecology clearly influenced single and co-infection probabilities. First, we found an effect of nesting behavior, with the probability to be single and co-infected being higher for species with open nests compared to closed nests. This result was also visible when considering each genus separately, though with some uncertainty regarding *Plasmodium*. Although contrasted findings have been reported (see e.g. table 1 in Ellis *et al.* 2020), several studies found a consistent support for a higher infection risk by haemosporidian parasites in species with open nests (e.g. Barrow *et al.* 2019; Ellis *et al.* 2020; Rodríguez-Hernández *et al.* 2021). This effect likely emerges because open nests can facilitate either the detection of cues used by vectors to locate their hosts (Yan *et al.* 2021) or the subsequent vector/host contact. Second, we found that the migratory behavior was an important predictor for both single and co-infection probabilities. The impact of migration on individual infection pattern provided contrasting results in the literature (e.g. Ricklefs *et al.* 2017; Poulin & Dutra 2021). Indeed, while migration may expose hosts to a broader range of parasites, resulting in higher parasite richness (e.g. Gutiérrez *et al.* 2019; Poulin & Dutra 2021) and higher infection risk (de Angeli Dutra *et al.* 2021), it may also be associated with lower infection prevalence by allowing individuals to escape environments presenting a high risk of infection or by culling sick individuals (Arriero & Møller 2008; Risely *et al.* 2018; Poulin & Dutra 2021). Our results are leaning toward the “migratory escape” and/or “culling” hypotheses; though the effect of migratory status was more uncertain when considering each parasite genera independently (possibly due to lower sample sizes).

Previous studies have identified the slow-fast life-history continuum as a robust predictor of different ecological patterns in multiple taxonomic groups (e.g. population dynamics, Marquez *et al.* 2019; success of invasive species, Ducatez & Shine 2019; range shifts Estrada *et al.* 2016). Our results add to this knowledge by showing that this continuum is also relevant to describe bird infection status by haemosporidian parasites. Indeed, although this factor was not a strong predictor of single infection probability, except for *Haemoproteus*, the co-infection probability was clearly lower for slow-living species compared to fast-living species. This result contradicts the widespread assumption that slow living species should display a higher infection probability because the likelihood to encounter parasites is more important owing to their long lifespan (e.g. Poulin & Morand 2000; Gutiérrez *et al.* 2019, but see e.g. Cooper *et al.* 2012). However, it has also been hypothesized that fast- and slow-living species are investing differently in their immune system and particularly in their adaptive (specific, less self‐damaging) immune response (Millet *et al.* 2007; Lee *et al.* 2008; Valenzuela‐Sánchez *et al.* 2021). For instance, to maximize their fitness along their lifespan, slow-living species may produce a more effective immune response (Tella *et al.* 2002; Millet *et al.* 2007) by producing secondary antibodies to a new antigen more rapidly, ultimately making it possible to control secondary infections more effectively (Elgert 2009) than fast-living species.

Haemosporidian co-infection probability also appeared to vary depending on the size of the host geographic range with larger co-infection probability for species having large distributions. This result is consistent with the “geographic range” hypothesis which postulates that widespread hosts face higher parasite pressure (Price *et al.* 1988). While only a handful of studies have examined the relationship between species geographical range and infection risk, yielding mixed results (Tella *et al.* 1999; Mlynarek *et al.* 2012; Suhonen *et al.* 2019), geographical range size has been shown to correlate positively with parasite richness in a wide range of diseases, from viruses and fungal parasites of plants (e.g. Mitchell & Power 2003; Miller 2012) to protozoan and metazoan parasites of birds and mammals (e.g. Lindenfors *et al.* 2007; Gutiérrez *et al.* 2019; Dáttilo *et al.* 2020). Host species with high parasite richness may have a higher probability of encountering several parasites during their lifetime, and ultimately of being co-infected, than host species with low parasite richness (but see Johnson & Hoverman 2012). Further, species with wide geographic ranges are usually assumed to have a broader range of tolerance to different ecological conditions (e.g. we found a positive relationship between species geographical range size and climatic niche breadth) and are thus considered as generalists (e.g. Slatyer *et al.* 2013). Generalist species, in addition to be more tolerant to resources and climate, may have also evolved a higher tolerance to the pathological effects of infection (Dennis *et al.* 2011; Barthel *et al.* 2014), which may result in an increase in the prevalence of co-infection in bird populations owing to a higher survival probability of each individual.

Finally, bird phylogeny explained a significant part of the variability in haemosporidian (co-)infection status, suggesting that host susceptibility, exposure or a combination of the two may be conserved across the time scale of avian diversification. Host–parasite associations usually tend to present a strong phylogenetic structure and host phylogeny has been shown to be a relevant predictor of disease spill-over and infection risk in a range of associations (e.g. plant-fungal pathogens, Gilbert & Webb 2007; amphibian-fungal pathogens, Greenberg *et al.* 2017; mammals-virus Streicker *et al.* 2010). However, previous studies usually found a low congruence between the phylogeny of haemosporidian parasites and the one of their avian hosts, partly due to host switching across large phylogenetic gaps (e.g. Ricklefs *et al.* 2014; Alcala *et al.* 2017). The extent to which susceptibility to haemosporidian infection is phylogenetically conserved rather than labile across bird phylogeny is therefore not well established (e.g. González *et al.* 2014; Barrow *et al.* 2019). While several factors explaining variation in single and co-infection probability tend to be phylogenetically conserved (nest structure, Fang *et al.* 2018; position on the slow-fast life-history continuum, Valenzuela‐Sánchez *et al.* 2021; geographic range size, Waldron 2007), we found support for a strong additional phylogenetic signal in infection probability. This result echoes with recent findings on Amazonian bird community (Barrow *et al.* 2019), and further highlights that host phylogeny is a much stronger determinant of co-infection than of single infection probability. Co-infections can therefore act as a stronger selective pressure than single infections that may ultimately influence on hosts evolution. The additional effect of host phylogeny observed here may be due to species differences in evolutionary conserved factors not included in our study such as the genetic component of the immune system (Minias *et al.* 2019) or other mechanisms allowing a higher tolerance to the pathogenic effect of parasite single and especially co-infection (Schneider & Ayres 2008; Sears *et al.* 2015).

Beyond the implications of this study for predicting host infection status, it demonstrates that tissue sample collections from museum specimens can be used to investigate evolutionary and ecological hypotheses at a relatively low cost and with no impact on wild populations. However, this also implies certain limitations. For example, our samples come from dead or injured individuals and our results may be biased towards weak individuals, who may have less resistance to pathogens (particularly for individuals found dead without injuries). In addition, we focused on haemosporidian parasites and we cannot rule out the possibility that birds have been infected by other blood parasites genus (e.g. *Trypanosoma*, *Babesia*) or by macroparasites such as helminths (e.g. Trematodes, Nematodes, Cestodes). We also concentrated on co-infections with different parasites genera and did not study mixed infections (involving different parasite lineages belonging to the same parasite genus). In order for future work to include mixed infections in their analyses, new specific PCR primers as well as more data (i.e. parasite lineages) in databases such as Malavi (Bensch et al. 2009) are needed to make effective use of computational phasing. Despite these limitations, our dataset and the associated results provided new insights regarding the role of species attributes and phylogeny in predicting bird (co-)infection status.

## CONCLUSION

By taking advantage of a large set of samples associated with museum specimens and using a state-of-the-art bayesian phylogenetic modelling framework, we identified relevant predictors of among-species differences in haemosporidian (co-) infection risks for western palearctic birds. Interestingly, we showed that our ability to predict co-infection risk was much higher than single infection risk, suggesting that random processes (e.g. encounter rate) may be more prevalent for the latter than for the former, where deterministic processes (e.g. species attributes, evolutionary history) are playing an important role. This result may explain why previous studies found no consistent patterns regarding the factors affecting single infection risks (see e.g. table 1 in Ellis *et al.* 2020). The strong phylogenetic signal that we found regarding co-infection risk further suggest that co-infections can act as a strong selective pressure that drive host’s evolution. These co-infections have rarely been considered in studies investigating the evolutionary ecology of host-parasite interactions and we therefore encourage future studies on this topic to do so. Expanding our understanding of the distribution of co-infection risk across multiple host species together with its effect on the evolution of both hosts and parasites will not only help us understand why some species are more susceptible to (co-)infection than other, but will also be of importance to address urgent public health problems regarding the emergence and evolution of infectious diseases.

## Authorship

RP, MC, PC and OG designed the study, MB, MA, AC, AK and OG collected data, RP and MC performed modeling work and analyzed output data, RP, MC and CSC wrote the manuscript with input from all authors.

## Data accessibility

Data supporting the results are archived in Dryad Digital Repository (https://doi.org/XXXXXXXXXXX) and are also provided in Supplemental Materials Appendix 1 to directly accompany this publication.

## Acknowledgements

We would like to thank Jérôme Wassef for his help in preparing the tissue samples. This project was funded by the Swiss National Science Foundation (SNSF), grants 31003A_159600 and 31003A_179378 to PC and CRSK-3_190197 to RP.

## Appendix 1

**Table S1**. **Table of avian ecology and life history traits.** This table summarizes the number of individuals per species, the infection prevalence per species but also includes species reproductive traits, maximum lifespan, behavioral traits (nest type, migration status), climatic niche breadth and position, the coordinates of each species on the PCA axes performed on birds reproductive traits and the coordinates on the axes of the Hill-Smith ordination.

**Table S2. Table of avian trophic niche traits**

## Appendix 2

**Figure S1. Analyses without full and facultative migrant species**. Owing to the difficulty to characterize the climatic niche of migratory species, the analyses presented in the main text (**Fig. 3**) were repeated with full and facultative migrant species excluded to test the robustness of our results. Analyses were conducted on 401 individuals of 45 species. The figures show the posterior mean and associated 95% credible intervals of predictors and random effects (i.e. phylogeny and host species) as estimated by *brms* models with infection status (i.e. uninfected, infected, co-infected) as the response variable. (**A**) *brms* model with climatic niche breadth and climatic niche position (KDE) fitted as predictors. (**B**) *brms* model with geographic range size fitted as predictor. Effects shown are relative to the uninfected reference category, with single infection in light blue and co-infection in dark blue. The posterior distribution of predictors and random effects with a negligible effect on infection and co-infection probabilities were expected to be centered on zero. For categorical predictors, the effects shown are relative to the reference categories: nest type (closed), migration behavior (resident), museum (Lausanne).

**Figure S2. Posterior mean and associated 95% credible intervals of the three different estimation of the climatic niches breadth and position as estimated by *brms* models with infection status (i.e. uninfected, infected, co-infected) as the response variable.** (**A**) Analyses carry out on the full data set and on the (**B**) data set where full and facultative migrant species were excluded. For each of the two data sets, three models were run. Only the method used to estimate the species realized niches was different. Indeed, in addition to kernel density estimators (KDE) presented in the main text we used two other algorithms to delineate species realized niches: convex hull (Chull) and alpha hull (Ahull). Chull are defined as the smallest convex set that contains all samples. Ahull and KDE are extensions of Chull where non-convex and irregular shapes are allowed. While Chull are sensitive to outliers, Ahull and KDE are sensitive to the density of points in the environmental space. Chull is parameter-free whereas Ahull and KDE depend on a parameter that controls the shape of the envelope. For the KDE, the bandwidth parameter was estimated from the data using a Hpi multivariate generalization of the plug-in bandwidth selector and species envelopes were defined as the minimum threshold of probability density that included 99% of points. For the Ahull, we adopted an iterative procedure staring with an alpha value of one and then incrementing the value of the parameter by steps of 0.2 until 99% of points were included in the envelope. From species envelopes, we extracted its area (niche breadth) and computed its centroid as the mean of the vertices. We then extracted the coordinates of the centroid on each of the two climatic PCA axes. Posterior mean estimates of geographic range size on infection and co-infection status was also plotted to the figure.

**Figure S3. Posterior mean and associated 95% credible intervals of the three different estimation of the climatic niches breadth and position as estimated by *brms* models with infection status (uninfected vs. infected) by each parasite genera as the response.** Posterior mean estimates of geographic range size on infection and co-infection status was also plotted to the figure.

**Figure S4. Haemosporidian infection across the avian phylogeny**. The proportion of individuals infected by each parasite genera for each well-sampled host species (≥ 5 individuals) was mapped as a continuous trait using the contMap() function in phytools (Revell 2012). The consensus tree was generated from 1000 phylogenies obtained from BirdTree.org (backbone tree from Hackett *et al.* 2008).

**Figure S5. Posterior mean and associated 95% credible intervals estimated by *brms* model with infection status (i.e. uninfected, infected, co-infected) fitted as the response variable and where (A) geographic range size or (B) OA1 was fitted as predictor instead of climatic niches parameters or slow-fast life history continuum respectively.** Effects shown are relative to the uninfected reference category, with single infection in light blue and co-infection in dark blue. The posterior distribution of predictors and random effects with a negligible effect on infection and co-infection probabilities were expected to be centered on zero. For categorical predictors, the effects shown are relative to the reference categories: nest type (closed), migration behavior (resident), museum (Lausanne).

**Figure S6. Posterior mean and associated 95% credible intervals estimated by *brms* models with infection status (i.e. uninfected vs. infected) by each parasite genera fitted as the response variable and where (A) geographic range size or (B) OA1 was fitted as predictor instead of climatic niches parameters or slow-fast life history continuum respectively.** Effects shown are relative to the uninfected reference category, with single infection in light blue and co-infection in dark blue. The posterior distribution of predictors and random effects with a negligible effect on infection and co-infection probabilities were expected to be centered on zero. For categorical predictors, the effects shown are relative to the reference categories: nest type (closed), migration behavior (resident), museum (Lausanne).

