## Appendix 2 for "Determinants of haemosporidian single and co-infections risks in western paleartic birds"

### Supplementary Material: Appendix 2

**Figure S1.** Analyses without full and facultative migrant species.

**Figure S2.** Posterior mean and associated 95% credible intervals of the three different estimation of the climatic niches breadth and position as estimated by *brms* models with infection status (i.e. uninfected, infected, co-infected) as the response variable.

**Figure S3.** Posterior mean and associated 95% credible intervals of the three different estimation of the climatic niches breadth and position as estimated by *brms* models with infection status (uninfected vs. infected) by each parasite genera as the response.

**Figure S4.** Haemosporidian infection across the avian phylogeny.

**Figure S5.** Posterior mean and associated 95% credible intervals estimated by *brms* model with infection status (i.e. uninfected, infected, co-infected) fitted as the response variable and where (A) geographic range size or (B) OA1 was fitted as predictor instead of climatic niches parameters or slow-Fast life history continuum respectively.

**Figure S6.** Posterior mean and associated 95% credible intervals estimated by *brms* models with infection status (i.e. uninfected vs. infected) by each parasite genera fitted as the response variable and where (A) geographic range size or (B) OA1 was fitted as predictor instead of climatic niches parameters or slow-Fast life history continuum respectively.

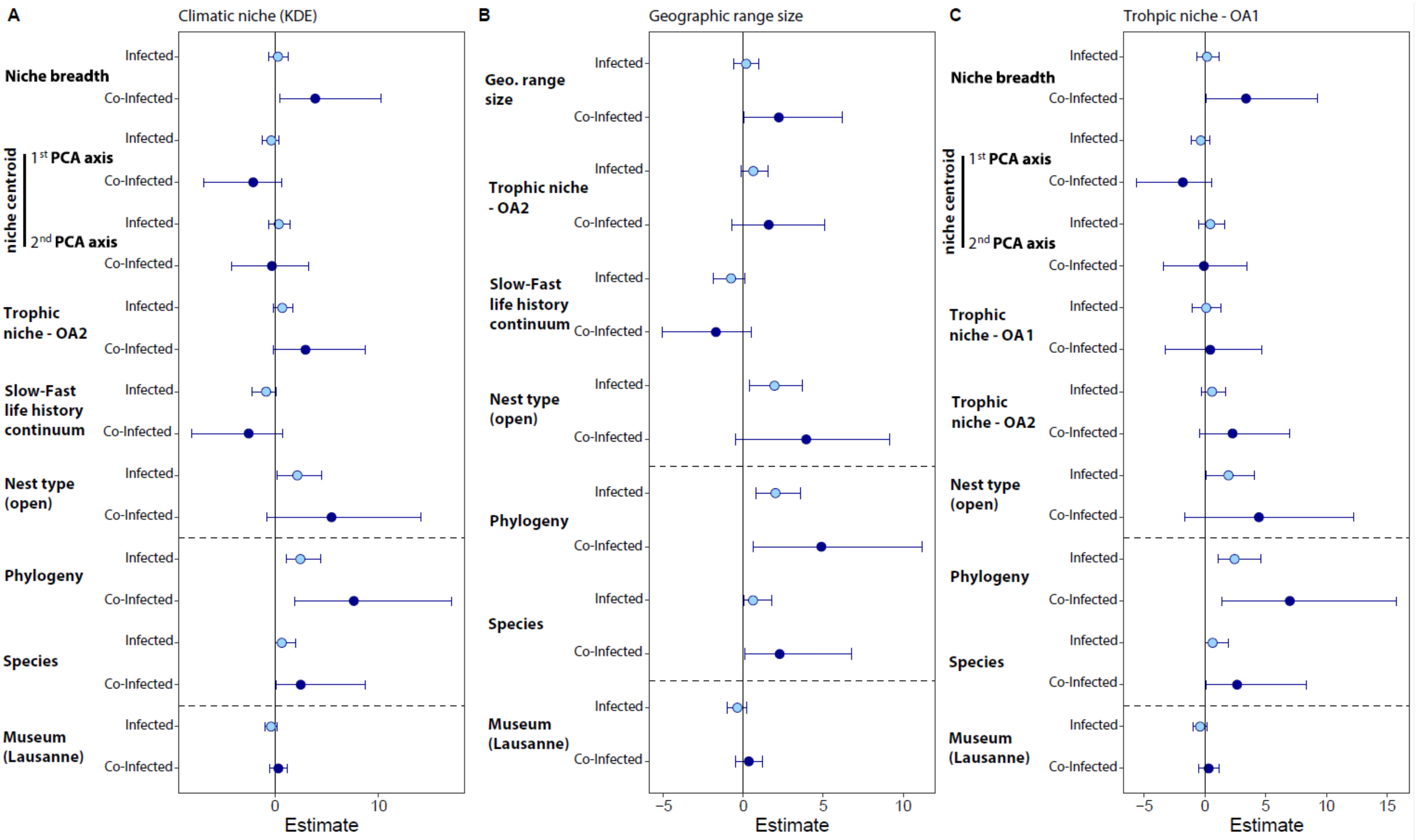


**Figure S1. Analyses without full and facultative migrant species**. Owing to the difficulty to characterize the climatic niche of migratory species, the analyses presented in the main text (**Fig. 3**) were repeated with full and facultative migrant species excluded to test the robustness of our results. Analyses were conducted on 401 individuals of 45 species. The figures show the posterior mean and associated 95% credible intervals of predictors and random effects (i.e. phylogeny and host species) as estimated by *brms* models with infection status (i.e. uninfected, infected, co-infected) as the response variable. (**A**) *brms* model with climatic niche breadth and climatic niche position (KDE) fitted as predictors. (**B**) *brms* model with geographic range size fitted as predictor. Effects shown are relative to the uninfected reference category, with single infection in light blue and co-infection in dark blue. The posterior distribution of predictors and random effects with a negligible effect on infection and co-infection probabilities were expected to be centered on zero. For categorical predictors, the effects shown are relative to the reference categories: Nest type (closed), migration behavior (resident), museum (Lausanne).

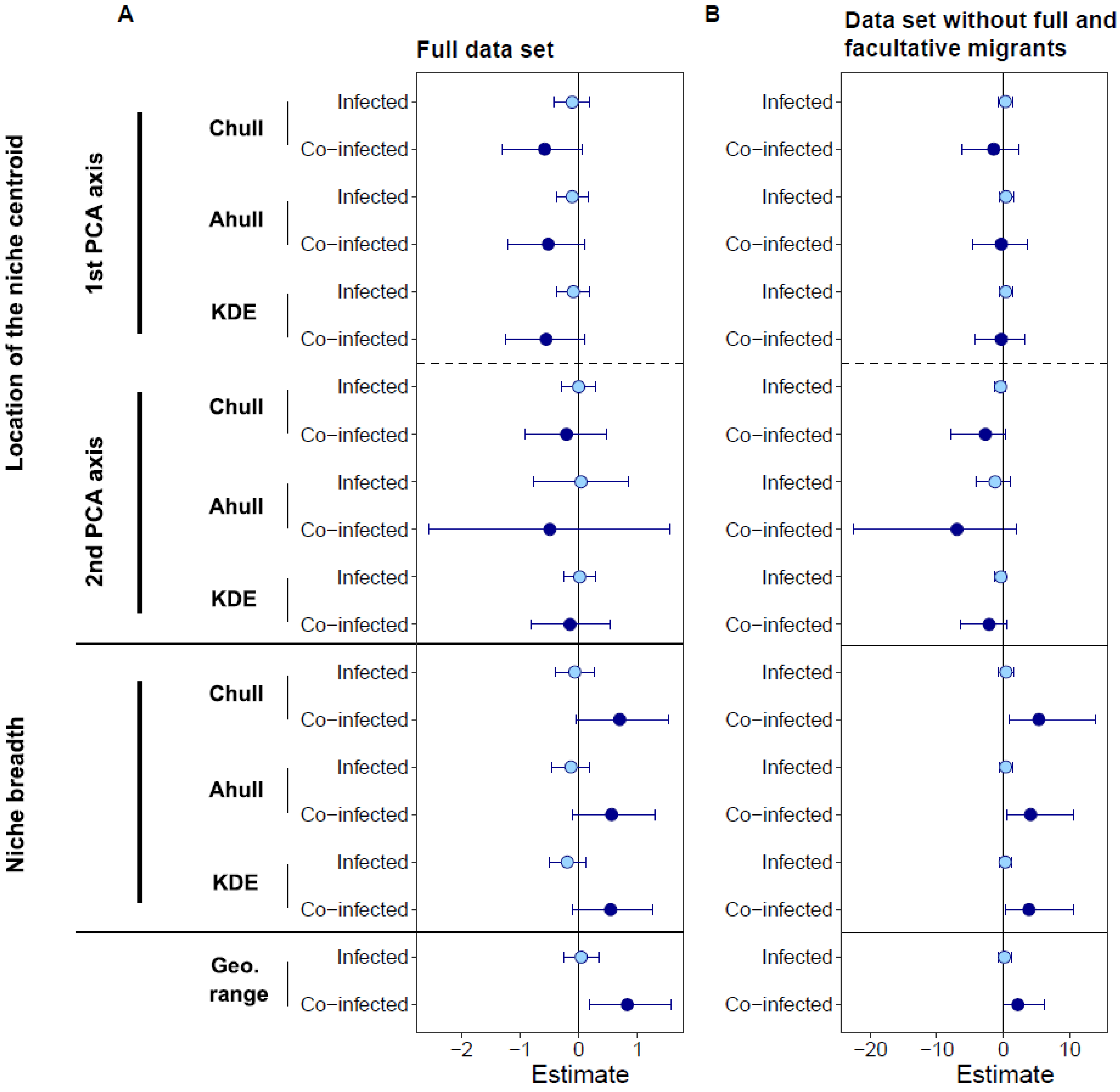


**Figure S2. Posterior mean and associated 95% credible intervals of the three different estimation of the climatic niches breadth and position as estimated by *brms* models with infection status (i.e. uninfected, infected, co-infected) as the response variable.** (**A**) Analyses carry out on the full data set and on the (**B**) data set where full and facultative migrant species were excluded. For each of the two data sets, three models were run. Only the method used to estimate the species realized niches was different. Indeed, in addition to kernel density estimators (KDE) presented in the main text we used two other algorithms to delineate species realized niches: convex hull (Chull) and alpha hull (Ahull). Chull are defined as the smallest convex set that contains all samples. Ahull and KDE are extensions of Chull where non-convex and irregular shapes are allowed. While Chull are sensitive to outliers, Ahull and KDE are sensitive to the density of points in the environmental space. Chull is parameter-free whereas Ahull and KDE depend on a parameter that controls the shape of the envelope. For the KDE, the bandwidth parameter was estimated from the data using a Hpi multivariate generalization of the plug-in bandwidth selector and species envelopes were defined as the minimum threshold of probability density that included 99% of points. For the Ahull, we adopted an iterative procedure staring with an alpha value of one and then incrementing the value of the parameter by steps of 0.2 until 99% of points were included in the envelope. From species envelopes, we extracted its area (niche breadth) and computed its centroid as the mean of the vertices. We then extracted the coordinates of the centroid on each of the two climatic PCA axes. Posterior mean estimates of geographic range size on infection and co-infection status was also plotted to the figure.

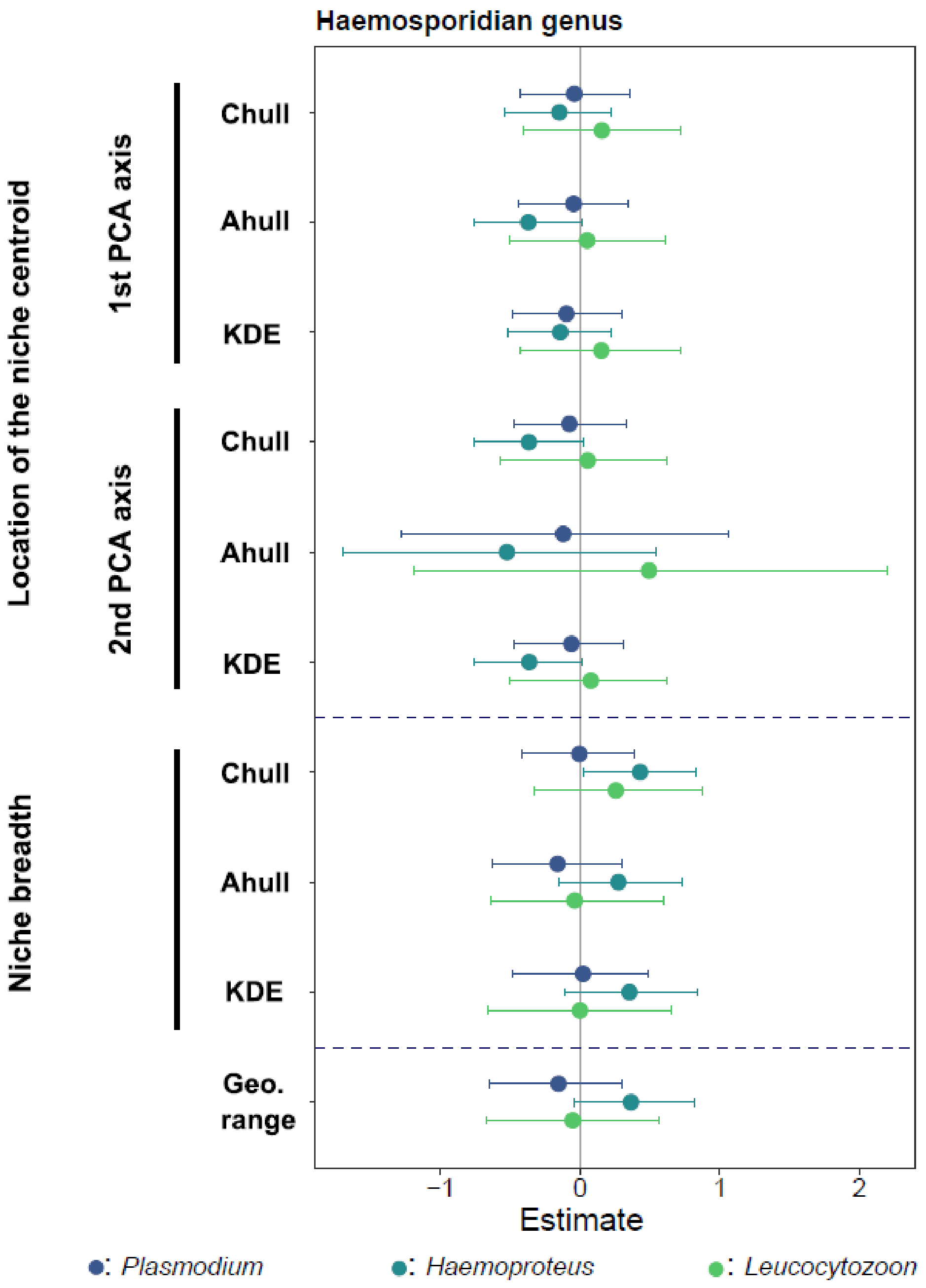


**Figure S3. Posterior mean and associated 95% credible intervals of the three different estimation of the climatic niches breadth and position as estimated by *brms* models with infection status (uninfected vs. infected) by each parasite genera as the response.** Posterior mean estimates of geographic range size on infection and co-infection status was also plotted to the figure.

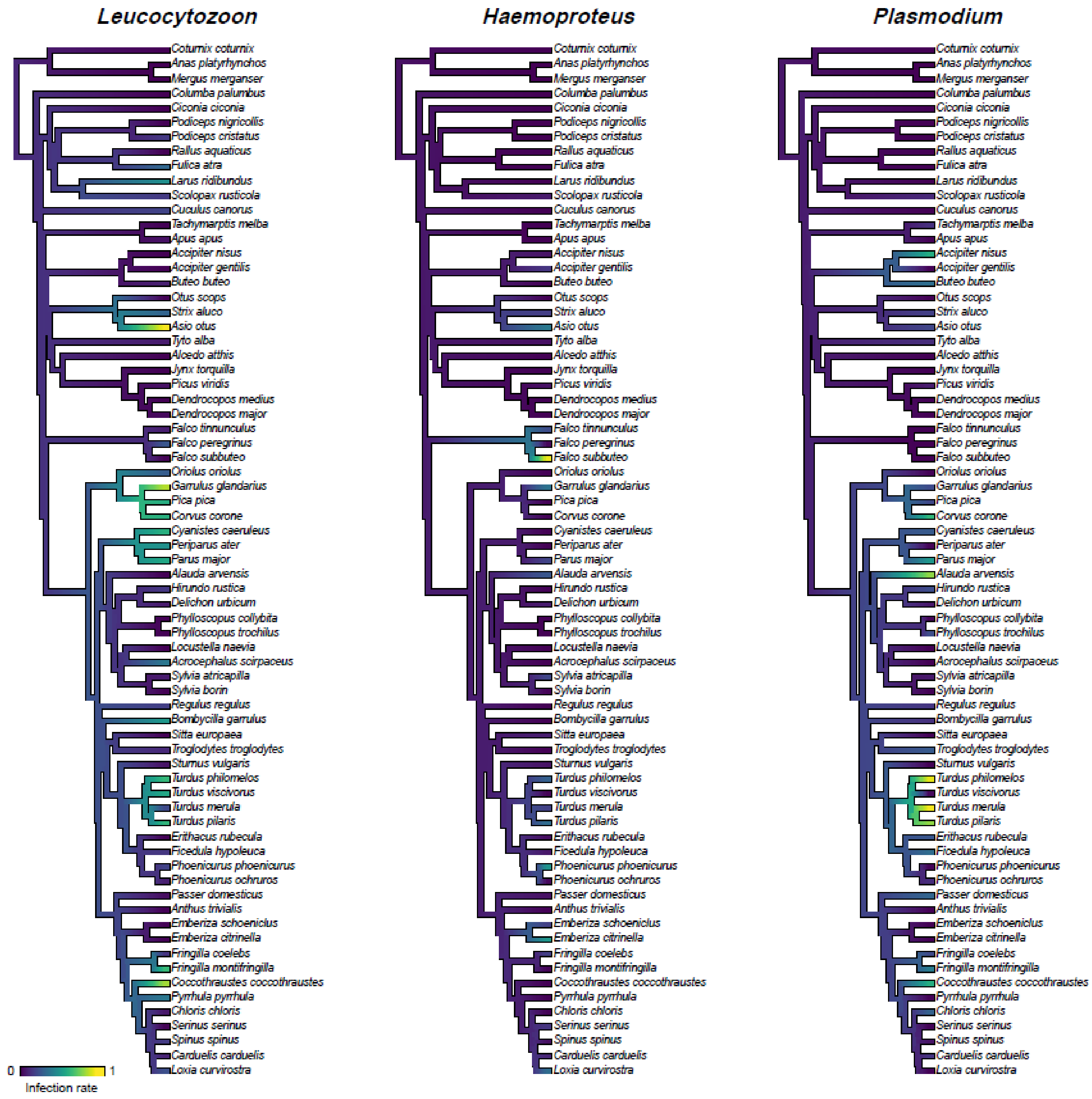


**Figure S4. Haemosporidian infection across the avian phylogeny**. The proportion of individuals infected by each parasite genera for each well-sampled host species (≥ 5 individuals) was mapped as a continuous trait using the contMap() function in phytools (Revell 2012). The consensus tree was generated from 1000 phylogenies obtained from BirdTree.org (backbone tree from Hackett *et al.* 2008).

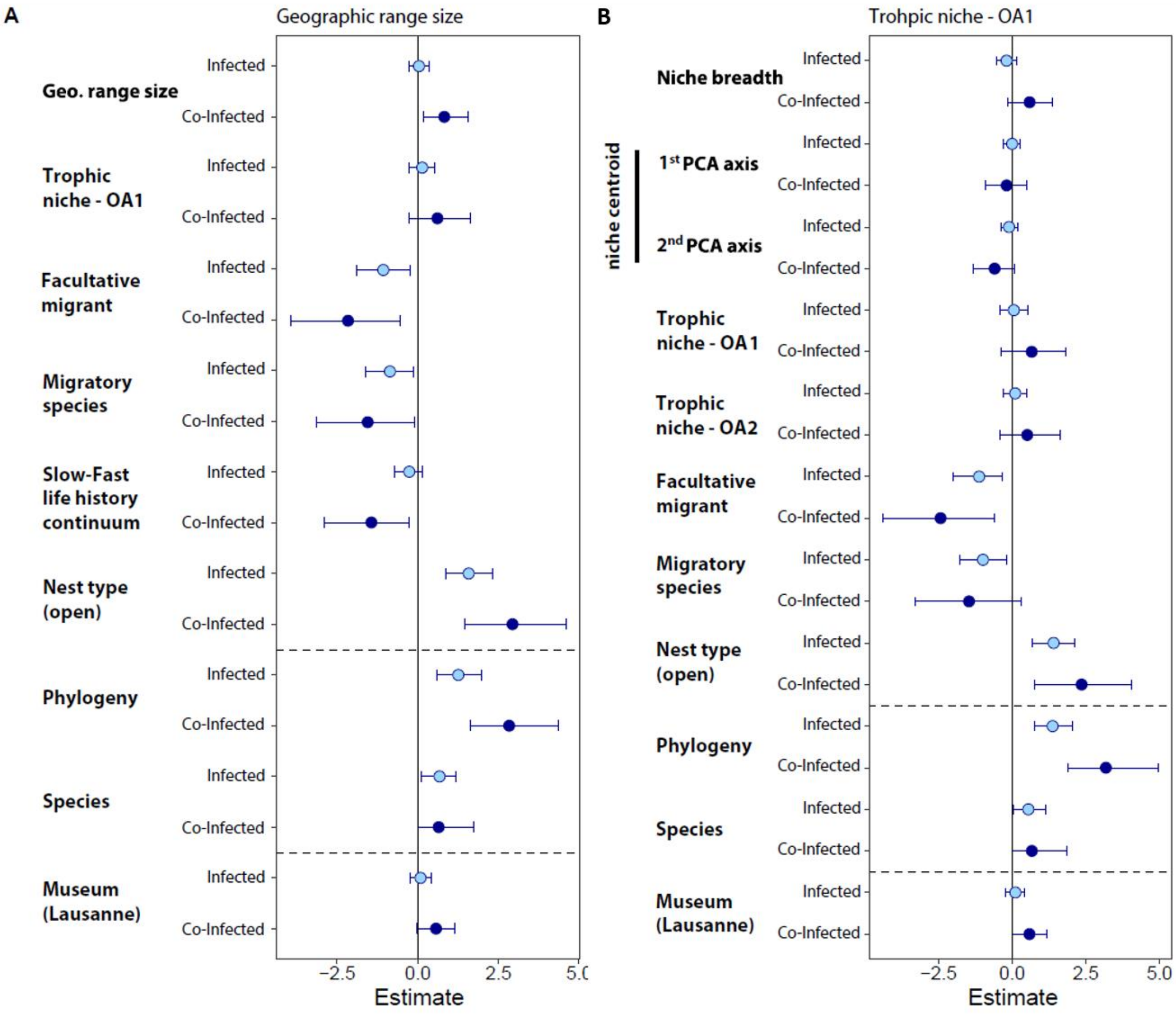


**Figure S5.** **Posterior mean and associated 95% credible intervals estimated by *brms* model with infection status (i.e. uninfected, infected, co-infected) fitted as the response variable and where (A) geographic range size or (B) OA1 was fitted as predictor instead of climatic niches parameters or slow-Fast life history continuum respectively.** Effects shown are relative to the uninfected reference category, with single infection in light blue and co-infection in dark blue. The posterior distribution of predictors and random effects with a negligible effect on infection and co-infection probabilities were expected to be centered on zero. For categorical predictors, the effects shown are relative to the reference categories: Nest type (closed), migration behavior (resident), museum (Lausanne).

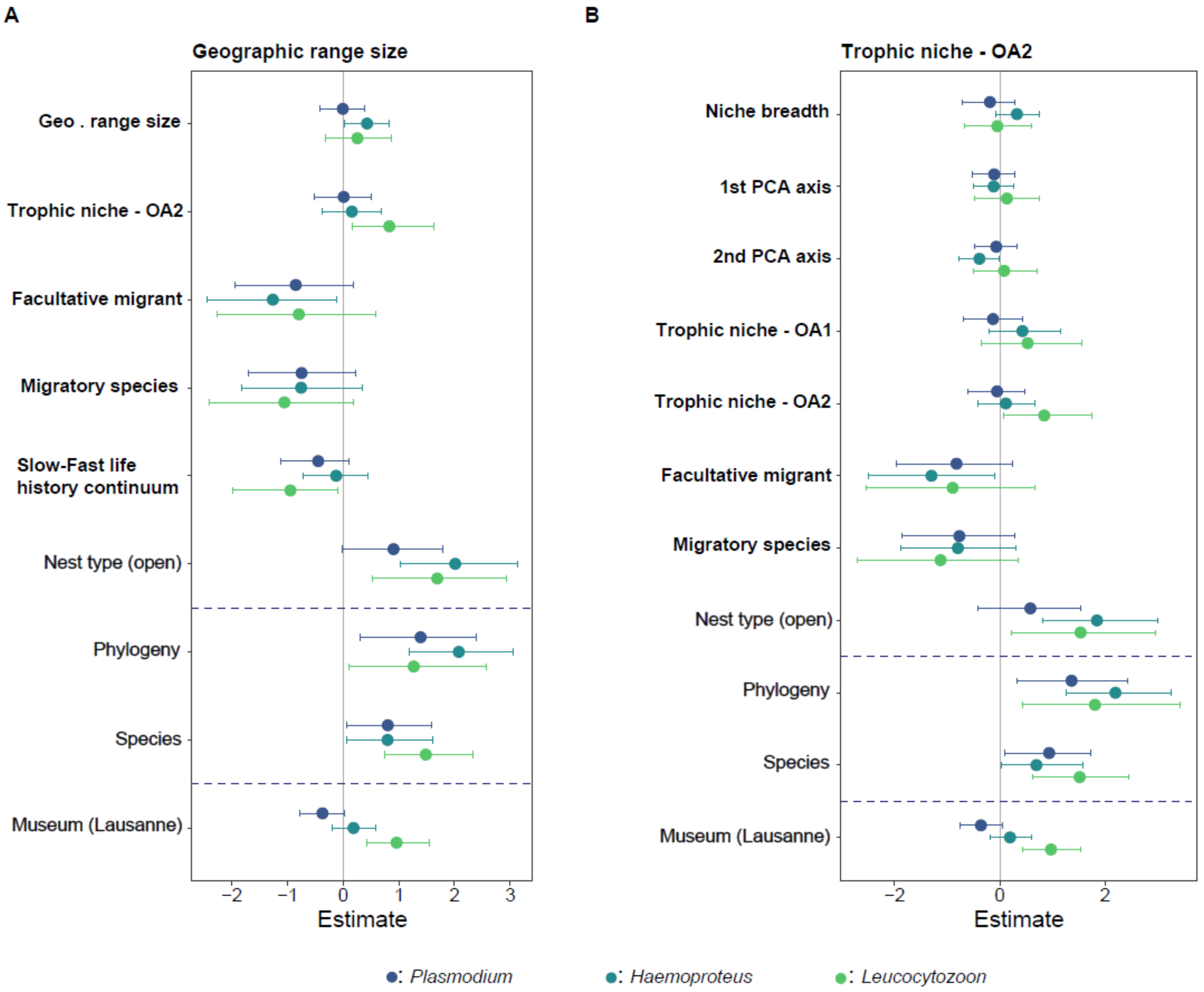


**Figure S6.** **Posterior mean and associated 95% credible intervals estimated by *brms* models with infection status (i.e. uninfected vs. infected) by each parasite genera fitted as the response variable and where (A) geographic range size or (B) OA1 was fitted as predictor instead of climatic niches parameters or slow-Fast life history continuum respectively.** Effects shown are relative to the uninfected reference category, with single infection in light blue and co-infection in dark blue. The posterior distribution of predictors and random effects with a negligible effect on infection and co-infection probabilities were expected to be centered on zero. For categorical predictors, the effects shown are relative to the reference categories: Nest type (closed), migration behavior (resident), museum (Lausanne).
